# Role of IL-6 and IL10 immune response genes against nematode *Haemonchus contortus* in ovis aries*- a novel report*

**DOI:** 10.1101/2023.08.30.555538

**Authors:** Kavita Rawat, Aruna Pal, Samiddha Banerjee, Abantika Pal, Paresh Nath Chatterjee, Subhasis Batabyal

## Abstract

*Haemonchus contortus* is one of the most deadly parasites affecting sheep industry causing heavy economic loss. In our earlier study, we have reported RIGI, an earlier established antiviral molecule and CD14 (an earlier established antibacterial molecule) to have potential role in antiparasitic immunity for the first time. In this current study, we aim to clone and sequence the IL-6 and IL10 genes in sheep (*Ovis aries*) and utilized bioinformatics methods to analyze and compare the functional and structural domains of peptides produced from IL6 and IL10. Furthermore, the study explored these domains in three-dimensional structures. The findings of this study provide molecular characterization of IL-6 and IL10 in sheep, which is essential for advancing our understanding of functional immune responses in this animal. We report IL6 and IL10 to have potential antiparasitic role against H. contortus for first time. The production of these molecules opens up possibilities for its potential use as recombinant proteins to manipulate the immune response. Evaluating its value in vaccine research and gaining a broader understanding of its role in immune responses becomes feasible with the availability of IL-6 and IL10. Moreover, the differential mRNA expression levels of IL-6, IL10 between healthy and diseased sheep indicate its crucial role in maintaining homeostasis and mounting immune responses. IL10 acts as receptor for H. contortus. The identification of this immune-responsive gene also offers new avenues for investigating long-term resistance to H. contortus infection in sheep.

## INTRODUCTION

T cells and macrophages secrete interleukin-6 (IL-6) to activate response after infection, trauma, burns, or other tissue damage after inflammation. A pleiotropic cytokine IL-6 with anti-inflammatory and pro-inflammatory effects that impact a number of different systems, including immunology, tissue repair, and metabolism **(Scheller *et al*., 2011).** Additional pro-inflammatory cytokines, such as TNF-, IL-1, and IL-6, promote the growth of B cells into plasma cells and activate cytotoxic T cells. Furthermore, it is an endogenous pyrogen that causes fever and the generation of acute-phase proteins from the liver **(Barkhausen *et al*., 2011).** After stimulation with the recombinant proteins, a numerous cytokine gene (IL-12p40, TNF α, IFN γ, IL-1, IL-4, IL-6, and iNOS) expression was evaluated in bovine peripheral blood mononuclear cells. All of the proteins stimulated the expression of IL-6, IL-12p40 and IFN γ but not of TNF α, IL-1, IL-4, or the iNOS gene. Except for TLR4, the expression of apoptosis-related genes remained unaffected.These findings demonstrate that B. abortus cellular antigens stimulate both cellular immunity and humoral in bPBMC via the production of IL-12p40, IL-6, and IFN γ without harming the cells **(Imet al., 2016).**Monophosphoryl lipid A and lipopolysaccharide, which are TLR4 agonists, increased the expression of some genes in sheep and human blood. Additionally, it was noted that the expression of the genes for key inflammatory mediators such as IL-1, IL-6, and IL-8, TNF alpha, NF-kappa B, PTGS2, PTX3, CXCL16, ETS2, CLEC4E, and KYNU was also increased**(Enkhbaatar *et al*., 2015).**Some data suggests that toll-like receptors 7 and 8 are involved in the induction of an antiviral response after SRLV (small ruminat lentivirus) infection, the production of IL-6, IFN, and TNF has been stimulated **(Blacklaws, 2012)**.

In a challenge experiment involving Haemonchus controtus, expression of several innate factors (SOD1 and GPX) as well as classical TH2 cytokines (IL-4, IL-5) was significantly increased (at 4 hours post-injury; 48 h). The expression of TH1 (IL-2 and IL-8) and Th2 (IL-4) cytokines was significantly higher in Pelibuey lambs on day 1 compared to day 0. Additionally, a positive correlation between the eosinophil count and the EPG count was found 48 hours after infection (P=0.05), indicating that lambs with more larvae developed have higher eosinophil activity.However, on day 14 PI, the pattern of gene expression was changed, and inflammatory cytokines like IL-6 and IL-8 were markedly down-regulated, but no additional alterations in gene expression were found**(Estrada-Reyes *et al*., 2015).**

The immune system of affected individuals is stimulated to reduce the damage caused by adult nematodes as well as prevent the development of third-stage GIN infective larvae (L3). The host immune system in response to the parasite invasion by inducing the generation of TH1 and TH2 cytokines (interleukin (IL)4, IL5, IL6, IL10, IL13, and others) in order to eradicate the infection caused by nematode**(Macrea *et al*.,2015).**In helminth infections and worm pathogenesis, IL5 and IL6 are the two cytokines that are thought to be most significant. In the TH2 response, IL5 is a critical modulator of the eosinophil arms, inducing eosinophil survival, differentiation, and chemotaxis, while IL6 is a cytokine with several functions that is related to the inflammatory response **(Naka *et al*., 2002).**

In a tropical region,changes in IL5 and IL6 gene expression in Pelibuey sheep can affect resistant and sensitive phenotypic characteristics to H. contortus, according to research on immune modulation methods identified against helminths**(Estrada-Reyes et al.,2015)**.T. circumcincta infection was found in the abomasal tissue of susceptible Blackface sheep and resistant Angus heifers that had been exposed to the Cooperiaoncophora and GINs Ostertagiaostertagi, research demonstrated greater production of proinflammatory cytokines such as Il-6**(Li et al., 2007 and Gossner et al., 2012).**In sheep that were naturally infected with Haemonchus species under two different nutritional conditions, the immunological response was characterised by significantly enhanced peripheral eosinophilia and an increase in blood concentrations of IgE, IgA, IgG, TNF β, and IL-6**(Toscan et al., 2017).**IL-6, an immune response gene, can be used in a variety of ways to treat illness. These include recombinant protein treatment, cloned insert gene therapy, and the evolution of disease-resistant transgenic or gene-edited animals.

Interleukin-10 is a cytokine with multiple, <u>pleiotropic</u>, effects in immunoregulation and inflammation, It also enhances B cell survival, proliferation, and antibody production.IL-10 is an immunosuppressive and anti inflammatory cytokine that regulates inflammation as well as Tcell, NK cell, and macrophages function (Kamnaka *et al*., 2006). It is mainly produced by TH2 cell but may also come from activated macrophages function. It is mainly produced by Th2 cells but may also come from activated macrophages. It targets Th1 cell, B cell, macrophages, NK cells, and mast cells. IL-10 selectively inhibits costimulation of Tcells by blocking CD28 phosphorylation.as a result its inhibit the synthesis of the Th1, cytokine IL-2, IFN-amma and TNF-alpha and oxidants by macrophages . It down regulates MHC class –II expression and stimulates production of IL-1ra. its receptor is CDw 210 and It also enhances B cell survival, proliferation, and antibody production (Moore *et al*., 2001).

In parasitic infection mainly Th1 and Th2 type immune response was associated (Finkelman and Urban, 1992; Janeway *et al*., 2004) and play important role in immunity against infection. IL-10 also inhibits the production of IFN-γ by Th1 and NK cells and induced the growth, differentiation and secretion of IgGs by B cells, which is responsible to eliminate the nematodes during infection, the host depend mainly on T lymphocytes, especially T helper 2 (TH2) cells .The TH2-type immune response induces the production of the various cytokines, such as IL-4, IL-5, IL-10, (Janeway *et al*., 2004) contributes to B cell differentiation leading to the expression of antibodies, such as IgE, IgG1, IgG4 and IgA and gathers eosinophils to target and eradicate the nematodes (Terefe *et al*., 2009). In studies it was obsereved that lambs that received *Taenia hydatigena* larvae vesicular concentrate prior to infection had a greater amount of eosinophils and mast cells and higher in situ expression of IFN gamma, IL-2, IL-4, IL-6 and IL-10 in the abomasal wall than the lambs that were infected with *H. contortus* only or that received ThLVC (Buendía-Jiménez *et al*., 2015).

Genome sequencing improves our ability to expand our basic understanding of associated species’ evolutionary paths. The present genome of each species has evolved over millions of years as a result of mutation and natural selection. The goal of this research is to clone and sequence the IL-6 gene in sheep (Ovis aries), a common household ruminant. Additionally, it emphasises using bioinformatics methods to recognise and contrast the functional and structural domains of peptides produced from IL6 as well as exploring the domains in 3D structures. To corroborate the aforesaid findings, in both healthy and diseased (Haemonchus infected) sheep, the IL-6 gene’s differential mRNA expression profile was examined.

## MATERIALS AND METHODS

### Collection of Samples and RNA Isolation

One gram liver tissue of sheep was acquired from a slaughterhouse run by the Kolkata Municipality Corporation. For the collection of samples, adult males (n=6) between the ages of 1-1.5 years were considered. For the purpose of isolating RNA, liver tissue was placed in a vial and soaked in Trizol before being delivered to the laboratory on ice. Total RNA was extracted according to protocol using the TRIZOL extraction method (Life Technologies, USA), and then used to synthesize cDNA **(Pal and Chatterjee, 2009; Pal et al., 2011).**Ethical approval was not required because the samples are collected from the slaughterhouse. The concentration of cDNA was calculated, and samples with concentrations of more than 1200 micrograms per ml were considered for further investigation. To carry out quantitative PCR expression profiling, abomassum tissue from both healthy and Haemonchus contortus-infected sheep was obtained. Both the tip and the middle of the abomassum were used to collect tissue. The expression from the abomasum’s tip was used as a control during the ddct computation.

### IL-6 and IL10 Gene PCR Amplification and cDNA Synthesis

20 μL of the reaction mixture contained 5 μg of total RNA, 10 mM of DTT, 0.5 μg of oligo dT primer (16-18mer), 1000 M of dNTP mix, 40 U of Ribonuclease inhibitor, and 5 U of MuMLV reverse transcriptase in a suitable buffer. The reaction mixture was completely blended and then incubated for one hour at 37°C. By heating the mixture unliganded for 10 minutes at 70°C, followed by cooling it on ice, the reaction might last up to 10 minutes. Following that, PCR was used to confirm the cDNA’s integrity. Nanodrop was used to determine the concentration of cDNA. The full-length open reading frame (ORF) of the IL6 and IL10 gene sequence needs to be amplified in sheep, a set of ovine primers was developed using DNASTAR software (Hitachi Miraibio Inc., USA).The primers, 80–100ng cDNA, 0.5μL of 10mM dNTP, 3.0μL 10X PCR assay buffer, 60ng of each primer,1U Taq DNA polymerase, and 2mM MgCl_2_made up the reaction mixture, which was contained in a volume of 25 microlitres. 35 cycles of initial denaturation at 94°C for 3 minutes, following denaturation at 94°C for 30 seconds, annealing at 61°C for 35 seconds, and extension at 72°C for 3 minutes were carried out on a thermocycler (PTC-200, MJ Research, USA). The final extension was carried out at 72°C for 10 minutes.

### Cloning and sequencing of cDNA

A 1% agarose gel electrophoresis was used to test the amplified product of the ovine IL-6 gene. A Gel extraction kit was used to purify the components from the gel (Qiagen GmbH, Hilden, Germany). A simple cloning vector called pGEM-T was used for cloning (Promega, Madison, WI, USA). After that, 10L of the ligated product was well mixed with 200L competent cells, and heat shock was delivered in a water bath for 45 secondsat 42°C. Following that, the cells were placed on cooled ice for 5 minutes before being introduced to the SOC medium.The centrifuged bacterial culture pellet was plated on an LB agar plate using 20 mg/mL X-Gal allowed for blue-white screening, Ampicillin (100 mg/mL) at a ratio of 1:1000, and 200 mg/mLIPTG. As described before, a small-scale alkaline lysis approach was used to isolate plasmids from overnight-grown culture **(Sambrook et al., 2001).**By using CD14 primers in PCR and restriction enzyme digestion, we characterised recombinant plasmids.The enzyme EcoRI(MBI Fermentas, USA) was used to generate IL-6 gene fragments and then added to a recombinant plasmid. Following that, these segments were sent through an automated sequencer using the dideoxy chain termination method with T7 and SP6 primers(ABI prism, Chromous Biotech, Bangalore).

### Sequence Analysis

The nucleotide sequence produced in this manner was examined for sequence alignments, protein translation, and contigs comparisons using DNASTAR Version 4.0, Inc., the USA.

### Study of Predicted sheep IL-6 and IL10 Protein Using Bioinformatics Tools

The predicted peptide sequence of the IL-6 and IL10 gene of CB cattle were developed using edit sequence (Lasergene Software, DNASTAR). Using Lasergene Software’s Megalign sequencing Programme (DNASTAR), this sequence was then aligned with the IL-6 and IL10 gene peptide of other livestock and avian species. The signal peptide of the IL-6 and IL10 gene was predicted using the tool (Signal P 3.0 Sewer-prediction results, Technical University of Denmark). A signal peptide is required for a cell to translocate a protein to the cellular membrane, and when the signal peptide is eventually cleaved, the mature protein is produced. In order to anticipate the presence and position of the IL-6 and IL10 gene’s signal peptide, a programme was utilised (Signal P 3.0 Sewer-prediction results, Technical University of Denmark). For protein stability and folding, disulfide linkages are crucial. The biologically active part of the protein is its three-dimensional structure.Utilising the proper software, di-sulfide bonds were predicted (http://bioinformatics.bc.edu/clotelab/DiANNA/)(Kim **et al., 2005).** Protein sequence-level analysis was employed (http://www.expasy.org/tools/blast/) to identify the locations of the GPI anchor and N-linked glycosylation sites.The molecule’s N-linked glycosylation determines whether it is membranous or soluble. In the case of membrane protein, a N acetylation site was discovered (MNSLFT), whereas the GPI anchor is responsible for anchoring. The presence of a C-mannosylation site has been discovered, and it plays a crucial biological function in signalling and cell-to-cell adhesion **(Loke *et al*., 2016).** O-linked glycosylation sites were located using the NetOGlyc 3.1 server (http://www.expassy.org/), and the NetNGlyc 1.0 programme (http://www.expassy.org/) was usedto evaluate Domain linker sites and Protein phosphorylation sites.Secondary Structure Predictions are used to predict the alpha helix and beta sheet,(Phyre 2).

### Prediction of three dimensions and assessment of model quality

To develop models and demonstrate the three-dimensional structure of the ovine IL-6 and IL10 gene, PyMOL (http://www.pymol.org/) was used.

### Protein-protein interaction network depiction

To understand the protein interaction network of the IL-6 protein, we ran a search in the STRING 9.1 database **(Franceschini et al., 2015).** Using a confidence score, the functional interaction was evaluated. Interactions with scores of less than 0.3, between 0.3 and 0.7, and greater than 0.7 are categorised as having low, medium, and high confidence, respectively. Additionally, we carried out a KEGG analysis to demonstrate the functional relationship between the IL-6, IL10 gene and other associated proteins.

### Molecular Docking analysis of IL6 and IL10 with *Haemonchus contortus* membrane protein

Patchdock and Firedock are two softwares used for molecular docking of IL6 and IL10 with the surface proteins of *Haemonchus contortus* as alpha tubulin and beta tubulin.

### IL-6 and IL10 gene differential mRNA expression profile between healthy and infected sheep

By counting the number of eggs per gramme, the sheep population used in this study was split into two groups: diseased (infested with the parasite *Haemonchus contortus*) and healthy (uninfected).

### Animals, faecal sample collection and FEC determination

At random, we took 60 sheep samples from the LFC farm, Mohanpur campus, WBUAFS. The samples were taken before the regular deworming treatment and they were taken prior to the usual deworming procedure and were thought to contain pre-existing gastro intestinal parasites in the previous three months. It was thought that the animals had come into contact with a Haemonchus infection while grazing.The sheep’s faeces were extensively analysed using the salt flotation method, and each sample had its Faecal Egg Count reviewed. Statistical analysis was performed to compare the outcomes of samples from two groups: healthy (xLJ + SD, where xLJ stands for mean and SD for standard deviation) and diseased (xLJ - SD, where xLJ stands for mean and SD for standard deviation). For the mutton-producing unit, sheep were routinely sold and slaughtered. The case study analysed a total of 12 tissue samples, of which 6 had high FEC (classified as diseased) and 6 had low FEC (identified as healthy). We took tissue samples from the lymph nodes, liver, abomasum, rumen, small intestine, and caecum.

### Blood sample collection

From 8 to 10 in the morning, a sheep’s jugular vein was aseptically punctured to obtain a blood sample with anticoagulants. 5ml sample was used to isolate DNA and stored at -20^0^C until used for analysis, while 2-3 ml of blood were obtained for haematological parameters, 5 ml of blood were collected for serum without anticoagulant, centrifuged for 10 min at 1000 rpm, and then stored at - 20 °C until analysis.

### Hematological Profiles

Haematological measures including packed cell volume (PCV), total leukocyte count (TLC), total erythrocyte count (TEC), haemoglobin concentration (Hb), E.S.R, and differential leukocyte count (DLC) were examined by standard methods described **(Jain et al., 1993).**

### Biochemical Analysis

The serum biochemical parameters were studied in the experiments such astotal protein, globulin,liver function test, albumin, albumin: globulin, ALP, ALT,AST,Indirect bilirubin,Total bilirubin, and direct bilirubin, and kidney function test, glucose,urea,uric acid, and BUNby using a semiautobiochemistry analyser (Span diagnostic Ltd.) with standard kits (Trans Asia Bio-Medicals Ltd., Solan, HP, India). The methodology used for estimation of ALP, ALT,total protein, total& direct bilirubin, albumin, glucose,creatinine urea, and uricacid were modified kind and king’s method,biuret method, 2-4-DNPH method, bromocresol green (BCG) method, GOD/PODmethod,modified Jaffe’s Kinetic method, GLDH-urease methodand trinder peroxidise method respectively.

### Real-Time PCR (qRT-PCR)

Each reaction was conducted on a 96-well optical plate of the ABI 7500 equipment using the same quantity of cDNA as determined by Nanodrop. Each reaction contained a 1ng cDNA template, 10 µl of 2X SYBR Green PCR Master Mix, 20 pMol of forward and reverse primers, and up to 20 µl of nuclease-free water. It was done in triplicate for each sample. Real-time PCR (qRT-PCR) was analysed using the delta-delta-Ct (ΔΔCt) method; Ct stands for the threshold value. Below is a list of the primers used in the QPCR investigation, each with an annealing temperature of 60^0^ C.

IL-6 primer: F-’TGC TGG TCT TCT GGA GTATC, R-GTG GCT GGA GTG GTT ATTG

IL-10 primer : F GCCACAGGCTGAGAACC, R-TTTCACTGCCTCCTCCAGAT

18S rRNA primer: F-TCCAGCCTTCCTTCCTGGGCAT, R-GGACAGCACCGTGTTGGCGTAGA.

## Result

### Molecular characterization of IL-6 and IL10 gene in sheep

Sequencing of IL-6 gene cDNA of the sheep resulted in the prediction of derived amino acids. According to reports, the IL-6 gene of sheep is 727 bp long and contains a peptide with a 208 amino acid sequence**(Figure 1)**.

**Figure 1:**
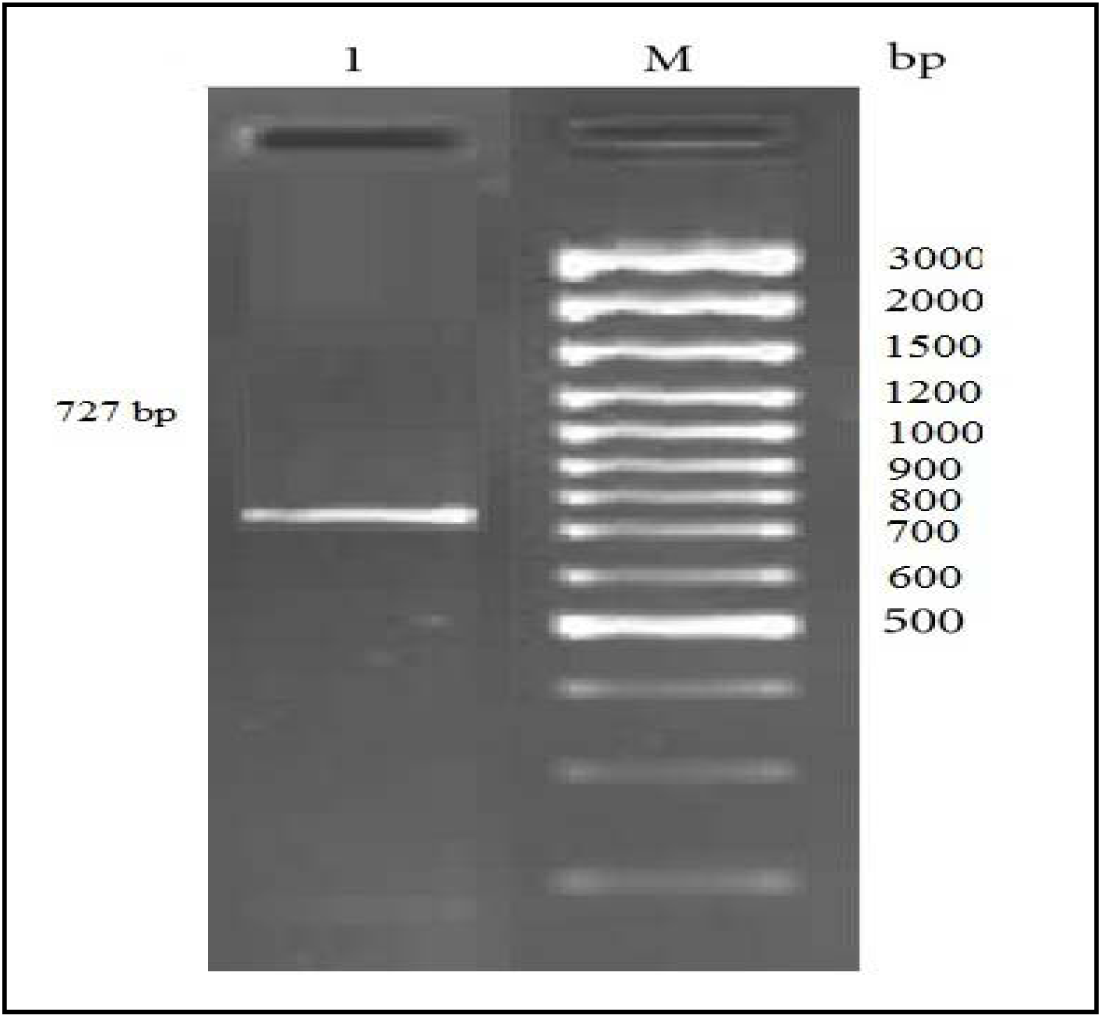
Lane 1: Amplification of 727 bp fragment of IL6-gene of by –PCR. Lane M: 100 bp DNA ladder plus.

### Illustration of the protein-protein interaction network and assessment of biological function

**Figure 4 shows**, using String analysis, how IL-6 interacts with other proteins in sheep. The interleukin-6 receptor and glycoprotein are in contact with the IL-6 sheep.There is a considerable interaction and functional overlap between the natural ligand for the neurokinin type 1 receptor (NK1R, a mediator of immune-modulatory activity), IL-6 as well as substance P (SP). The gp130 and IL-6R proteins create a complex when IL-6 binds with its receptor, activating the receptor. Signal transducers, Janus kinases (JAKs), and Activators of Transcription can be used to initiate a signal transduction cascade, these complexes combine the intracellular domains of gp130 (IL6ST) (STATs).

**Figure 2:**
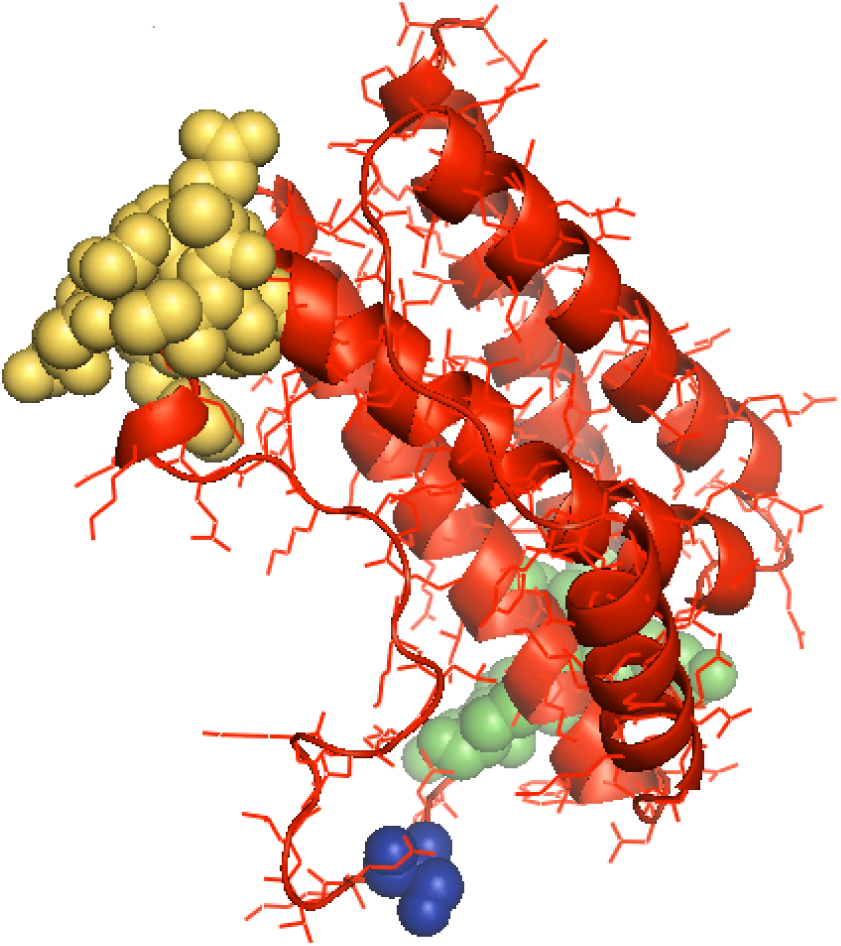
3D structure of ovine IL6 (red), **Di sulphide bond (aa position 72-78): green sphere** **Di sulphide bond (aa position 101-111): yellow sphere** **Modified residue Phosphoserine(aa position 81): blue sphere**

**Figure 3: :**
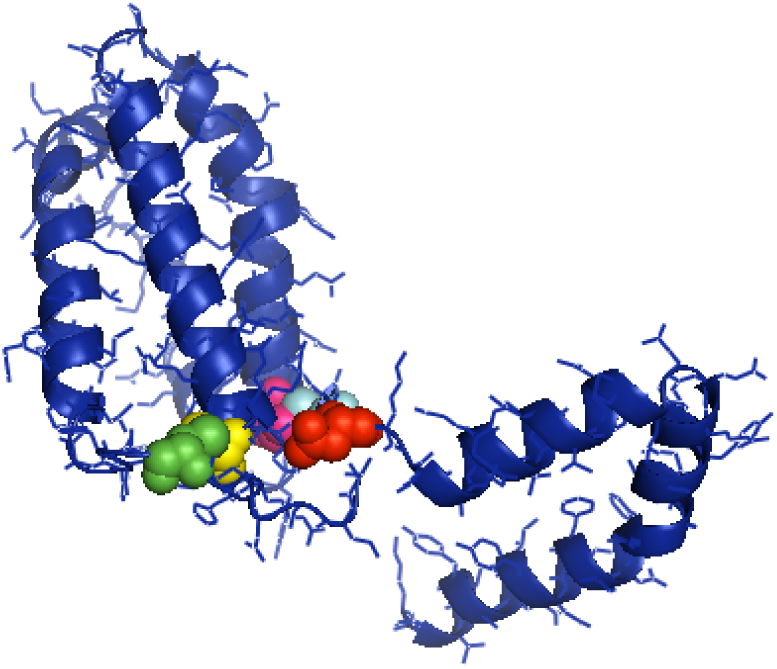
3D structure of ovine IL10 (blue), **Di sulphide bond(31-127)** Di sulphide bond (aa position31): green sphere Di sulphide bond (aa position 127): yellow sphere **Di sulphide bond(81-133)** Di sulphide bond (aa position 81): hot pink sphere Di sulphide bond (aa position 133): cyan sphere **N linked glycosylation (aa position 135): red sphere**

**Figure 2A:**
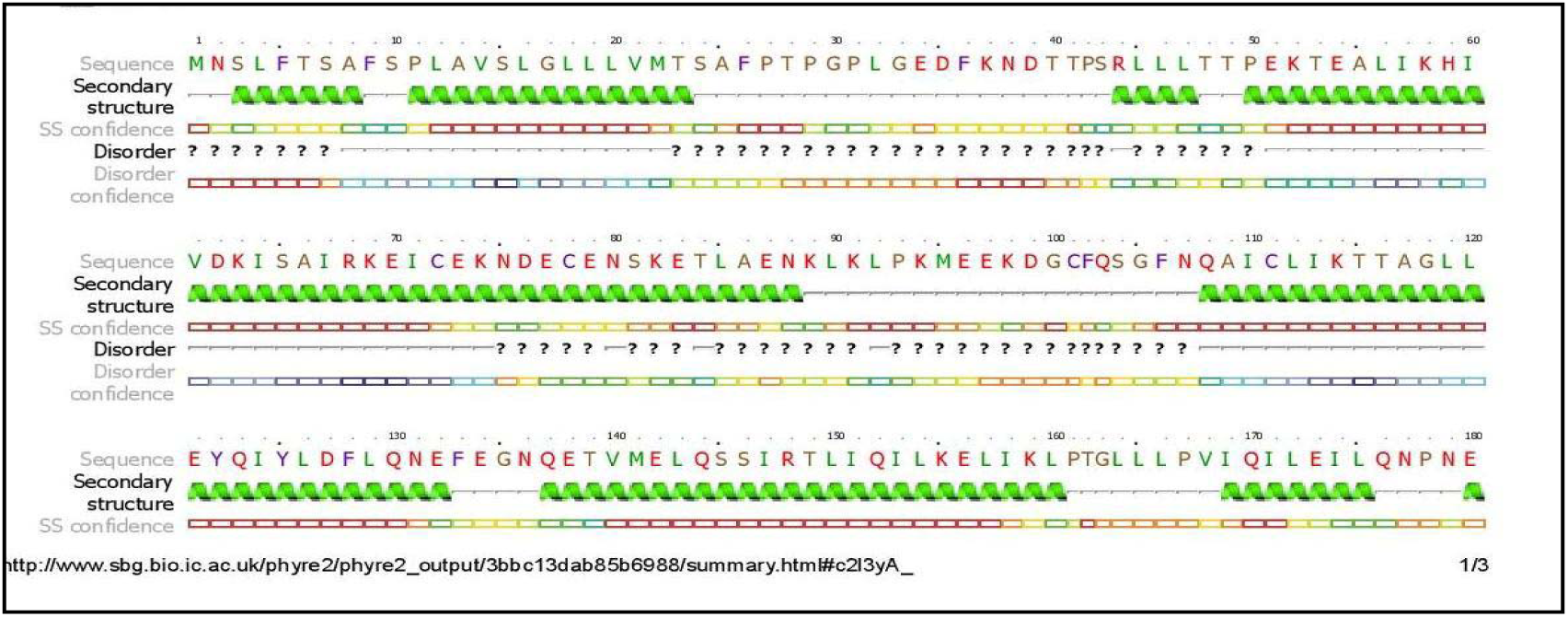
Secondary structure prediction of IL-6 gene.

**Figure 2B:**
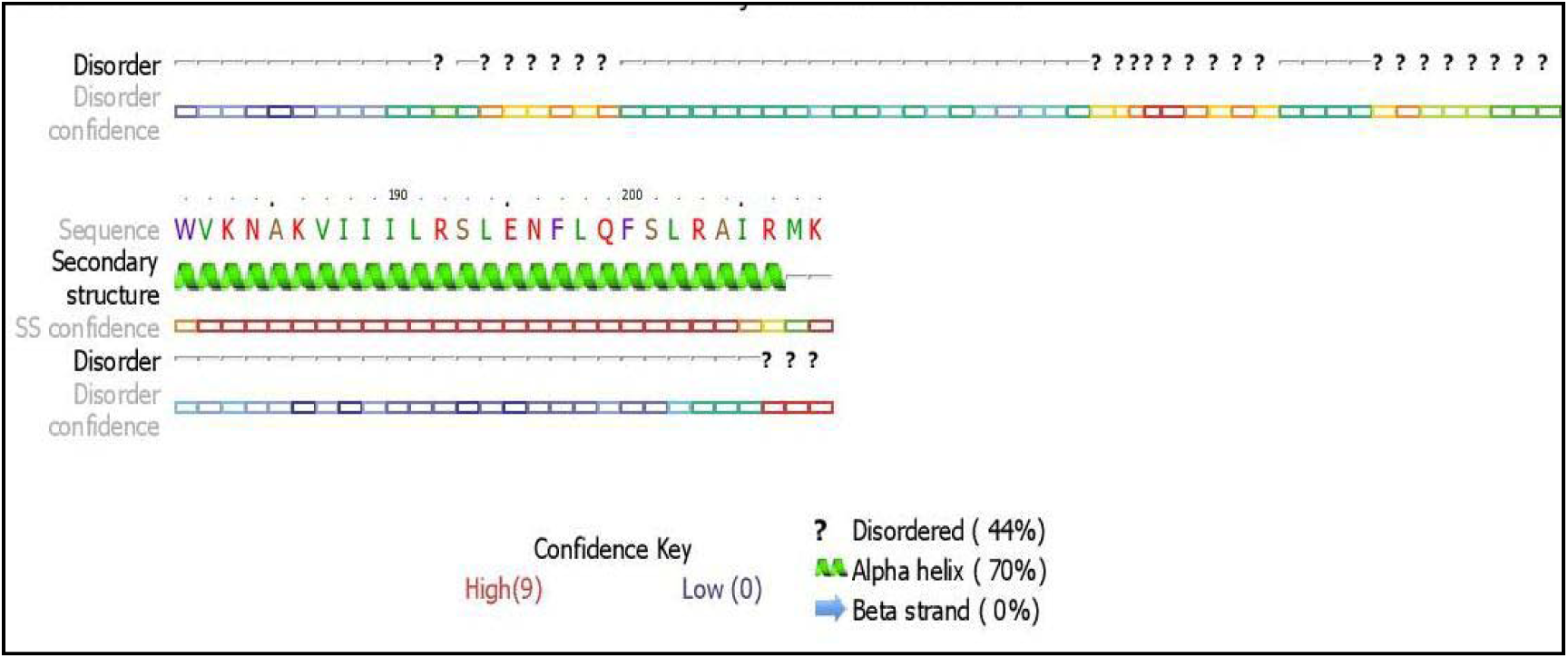
Protein Disorder predictionof IL-6 gene.

**Figure 3:**
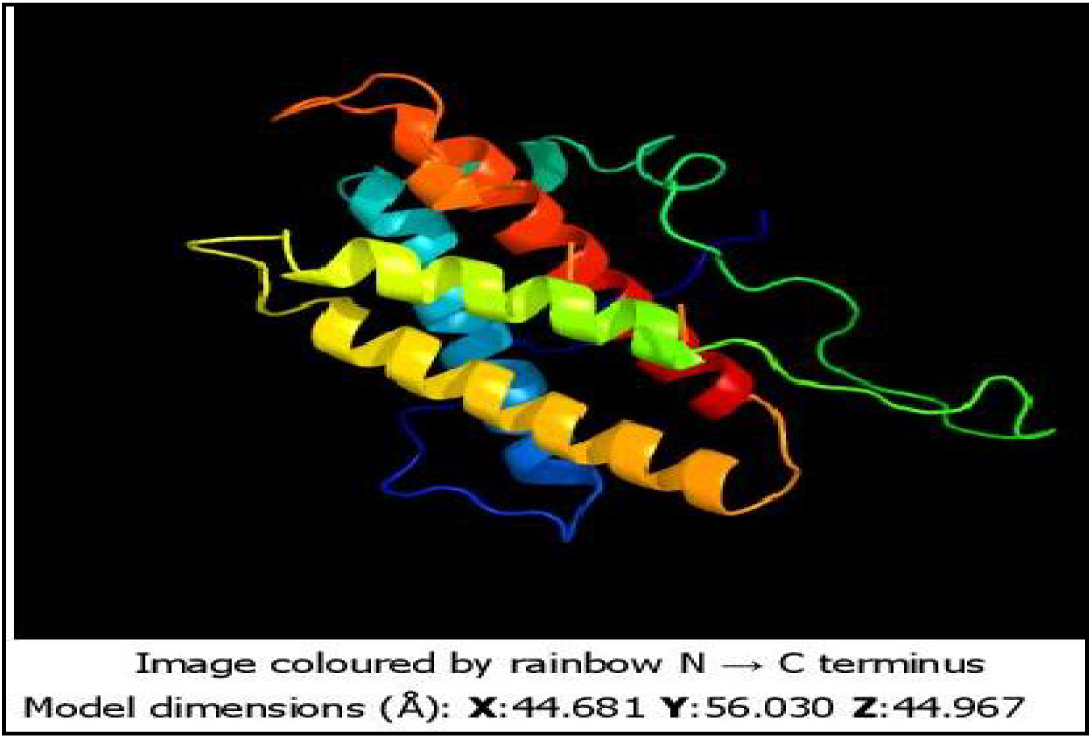
3-D Structure of IL-6 Gene through Pymol with secondary structure, depicted as Helical structure of IL-6 Gene.

**Figure 4:**
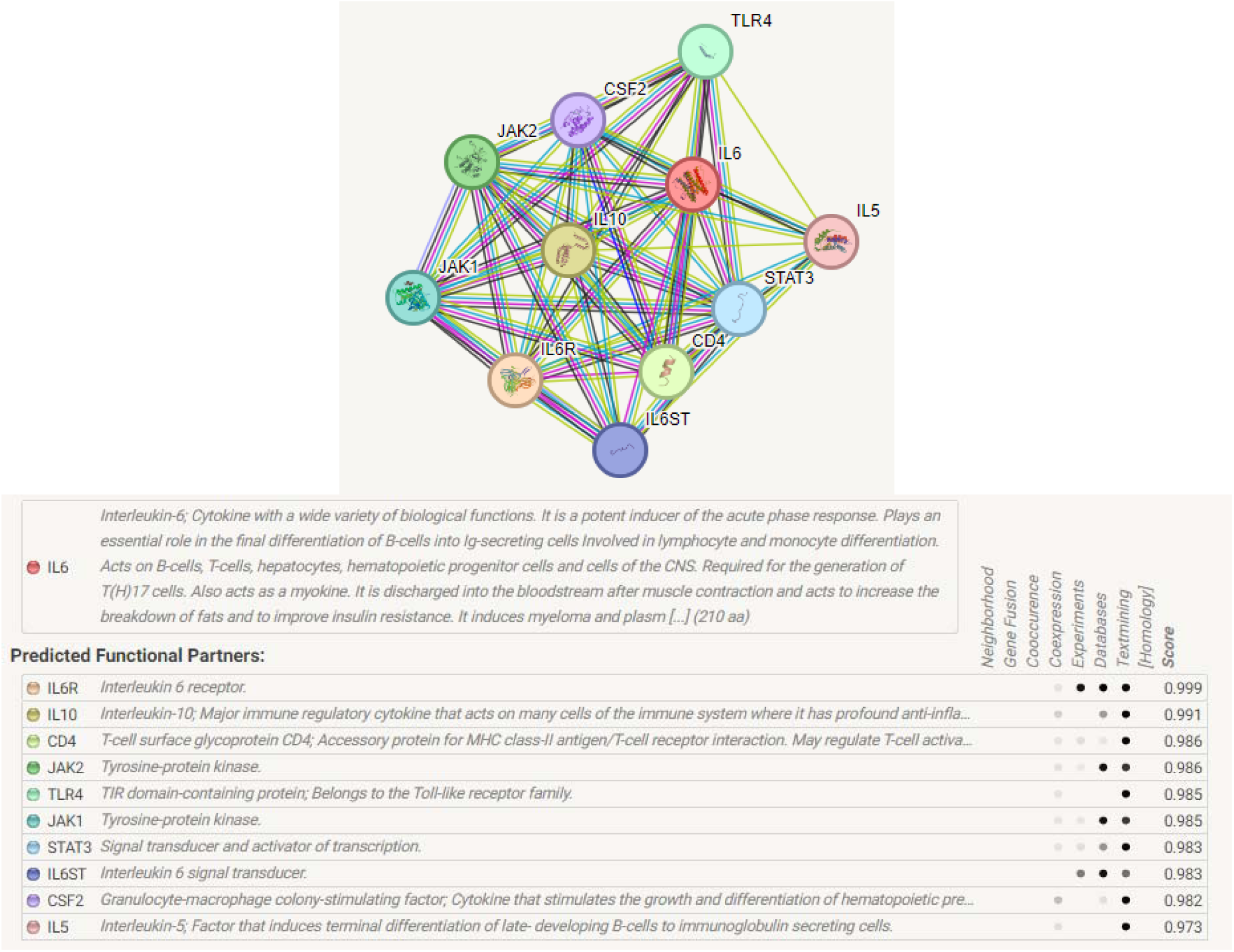
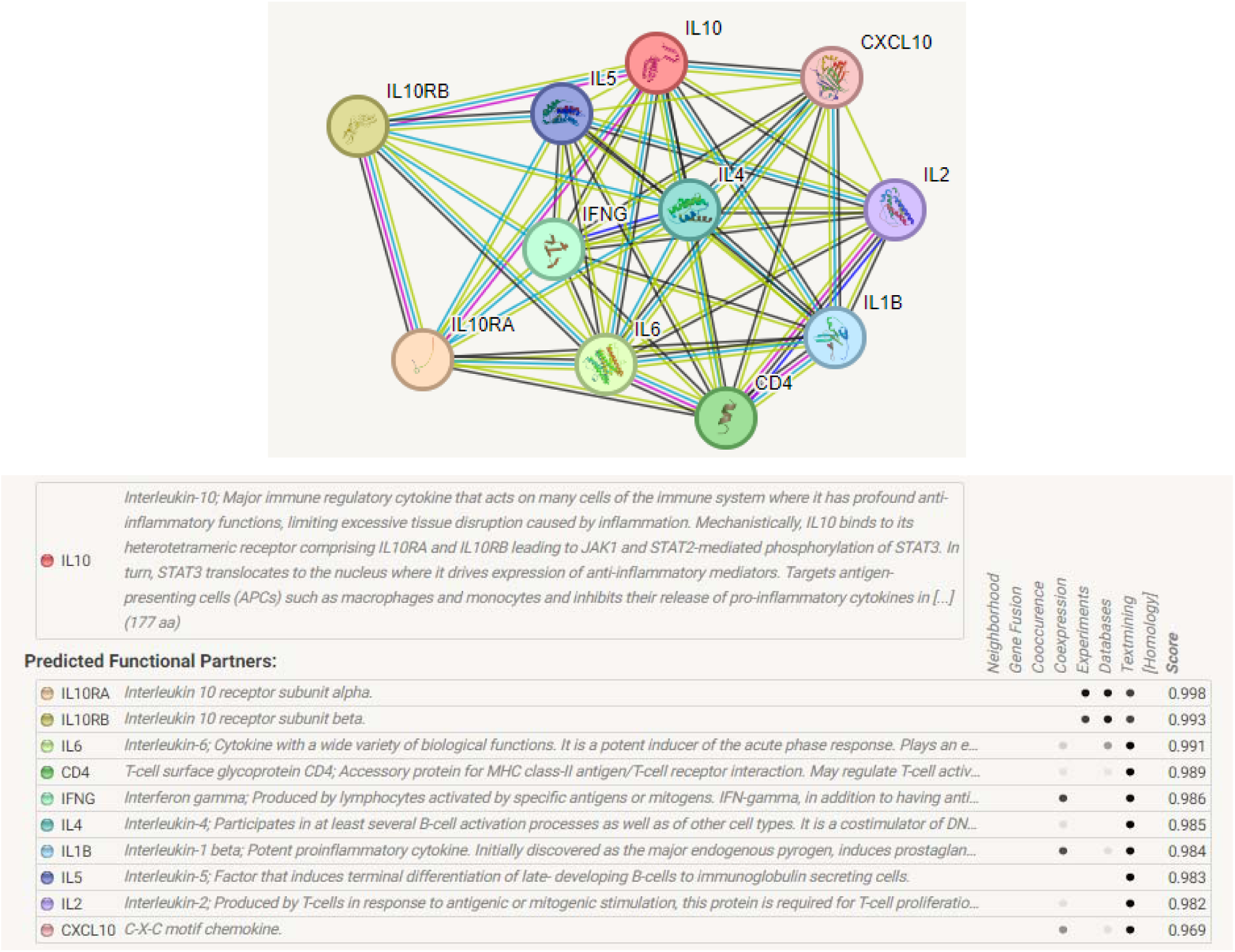
Protein-Protein Interaction Network of IL-6 Gene.

In our current study, mRNA expression level of IL-10 gene in disease sheep was high as compared to healthy as shown in fig.11.

IL-10 is a cytokine with multiple, <u>pleiotropic</u>, effects in immunoregulation and inflammation, It also enhances B cell survival, proliferation, and antibody production.IL-10 is an immunosuppressive and anti inflammatory cytokine that regulates inflammation as well as Tcell, NK cell, and macrophages function. (kamnaka *et al*., 2006). It is mainly produce by TH2 cell but may also come from activated macrophages function. It is mainly produced by Th2 cells but may also come from activated macrophages . It target are Th1 cell, B cell, macrophages, NK cells, and mast cells. IL-10 selectively inhibits costimulation of Tcells by blockin CD28 phosphorylation.as a result its inhibit the synthesis of the Th1, cytokine, IL-2, IFN-amma and TNF-alpha and oxidants by macrophages . It downreulates MHC class –II expression and stimulates production of IL-1ra. its receptor is CDw 210 and It also enhances B cell survival, proliferation, and antibody production (Moore *etal*.,2001).

In our present study IL-10 is overexpressed in disease animal compared to healthy one. In accordance with our study, over expression of IL-10 was also observed in the resistant breed Canary Hair Breed comparative to Canaria Sheep (Guo *etal.*, 2016).They also reported high induction of TH2 type immune response in *haemonchus controtus* infection.

Similarly some more studies (Gadahi *et al*., 2016) reported that increased expression of IL-10 in *in vitro* studies of goat’s PBMCs (peripheral blood mononuclear cell) challenged with *Haemonchus controtus* excretory and secretary products. Shakaya (2007) also revealed significant increased in IL-10 expression in *haemonchus controtus* infected Sufflock breed. Other Studies also demonstrated that *haemonchos controtus* infected Lambs that previously received Taenia hydatigena larvae vesicular concentrate (ThLVC) had a high eosinophils and mast cells and higher in situ expression of IL-10 in the abomasal wall compared to the lambs that were infected only with *H. contortus* and previously received ThLVC(Buendía-Jiménez *et al*., 2015).

However, Some studies reported high expression of the Th1-related cytokine genes (IL-1β, TNF-α, and IFN-γ)) day 3 and drastically up-regulated expression of Th2-related cytokine genes (IL-10 and IL-13) on day 7 in abomasal tissue of sheep after infection with *H. contortus* (<u>Hassan</u> *et al*., 2011). Further, evidences supported that *H.controtus* infection causes strong and early expression of TH1 and TH2 cells at 4 hours2 days post infection and elevated expression of IL-10 and FCεR1A, which may be involved in the immune regulation against nematode infection (Estrada-Reyes *et al*., 2017).

Hence, in parasite infection, antigen enter into the host cell, it is presented by APC through the MHC-II and it induced the secretion of cytokines from T cell, these cytokines stimulate immune response and activate lymphocyte, proliferation and differentation (Budhia *et al*., 2006). The Th1 and Th2 cell released different kinds of cytokines such as interleukin (IL-2), interferon-gamma (IFN-γ) and tumour necrosis factor-alpha (TNF-α) from TH1 cells and IL-10 produce from Th2 type cells and these cytokines are responsible to stimulates B cell differentiation, production of different immunoglobulin such as IgE, IgG1, IgG4 and IgA, mastocytosis, and eosinophil activation and function (Janeway *et al*., 2004) .Evidences also favours that Th2 cell mechanism play important role against *haemonchus controtus* infection (Shakya *et al*.,2011) Activated eosinophils are responsible to eliminate parasites (Terefe *et al*., 2009). .

In our present study the over expression of IL-10 may be due to Th2 response as it is associated with the production of IL-10 leading to eosinophil mobilization, intestinal mast cell accumulation and production of IgE, mucosal mast cell infiltration, intestinal eosinophilia, elevated serum IgE, increased level of parasite specific IgG-1 which are responsible to eradicate worms (Tizard, 2004).

### KEGG analysis

IL-6 works via several metabolic channels. The route for parasite resistance has been shown in **Figure 5** since immunization against parasitic infection is significant in the current investigation.

**Figure 5:**
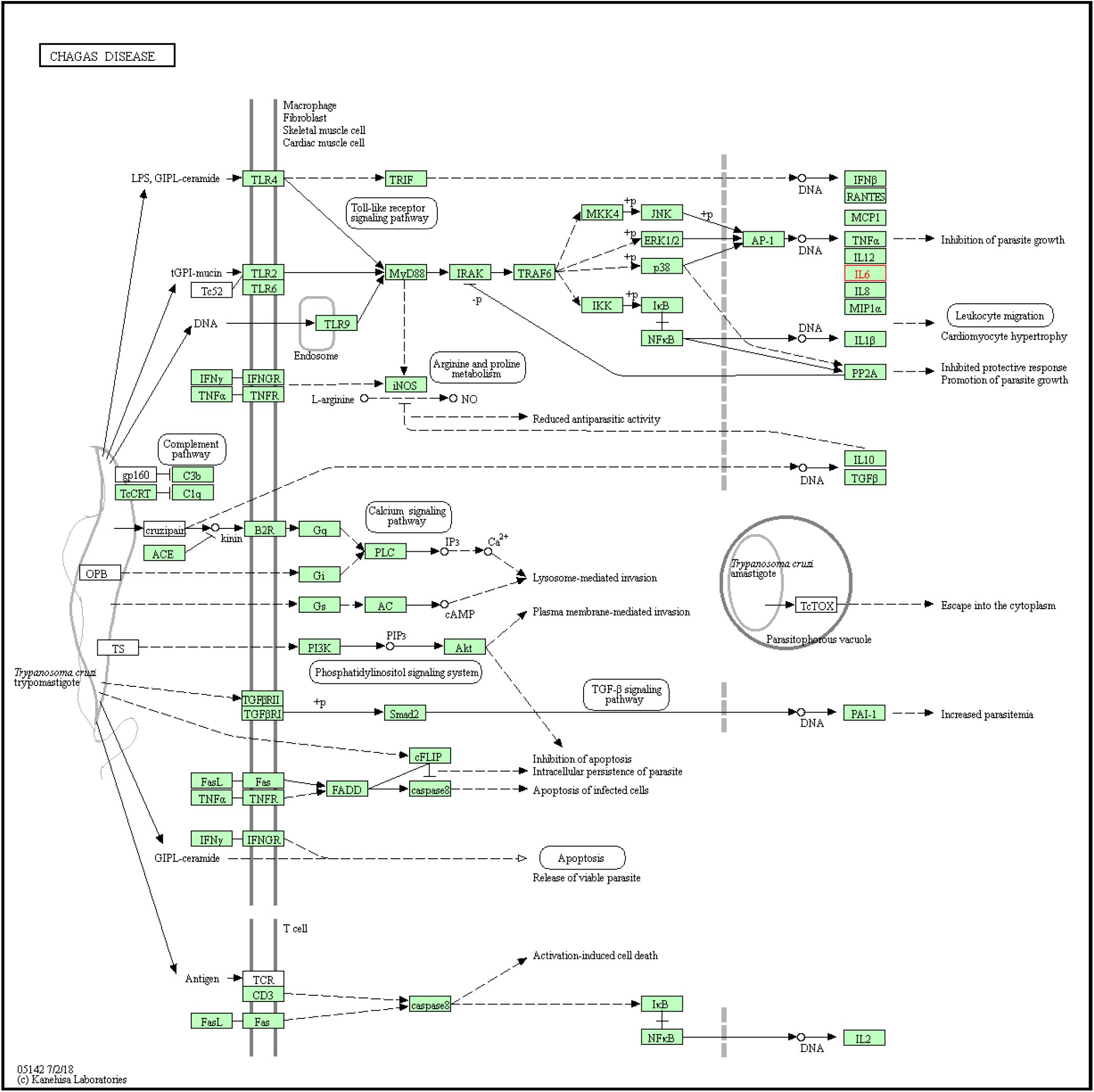
Pathway for Parasitic Resistance of IL-6.

### Molecular Docking of IL6 and IL10 with *Haemonchus contortus*

The patchdock score for IL6 with *Haemonchus contortus* alpha tubulin was observed to be 15210, whereas that of betatubulin was 14368. These results indicate low binding score, which means IL6 is not working as a receptor for binding to H. contortus.

However, higher patchdock score was observed for IL10 in comparison to IL6. Patchdock score was observed to be 18440 for beta tubulin, 18898 for alpha tubulin.These results indicate the role of IL10 as receptor and binding with H. contortus.

### Differential mRNA expression profiling for IL6 and IL10 genes Interleukin-6(IL-6) gene

#### mRNA Expression

As indicated in **Figure 6**, healthy sheep in our present study had higher levels of IL-6 gene mRNA expression than diseased animals. T cells and macrophages release IL-6 to promote an immune response after infection, following trauma, burns, or other tissue damage following inflammation.

**Figure 6:**
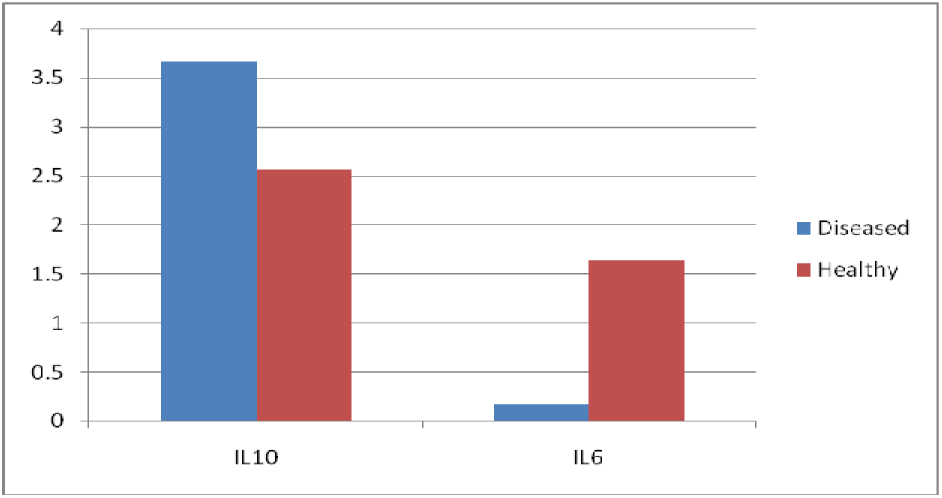
Differential mRNA expression profile of sheep immune response genes (IL10 and IL6) with respect to diseased and healthy conditions of parasitic infestation. *(Haemonchus controtus)*

#### Hematological and Biochemical parameters with respect to healthy and infected sheep

The haematological parameters were compared between sheep that were healthy and those that were infected with H. contortus (Table 1).

**Table 1:** Hematological parameters in diseased and healthy sheep.

| S.no | Parameter | Diseased | Healthy |
| --- | --- | --- | --- |
| 1. | Hb gm/dl | 7.96<sup>b</sup>±0.19 | 10.37<sup>a</sup>±0.19 |
| 2. | ESR mm | 1.00±0.00 | .90±0.04 |
| 3. | PCV% | 27.84<sup>b</sup>±0.67 | 36.25<sup>a</sup>±0.64 |
| 4. | TEC /cumm | 3.98<sup>b</sup>±0.05 x10<sup>6</sup> | 4.95<sup>a</sup>±0.08x10<sup>6</sup> |
| 5. | TLC/cumm | 9200.00<sup>a</sup>±159.23 | 6587.50<sup>b</sup>±210.81 |
| 5. | Neutrophil % | 53.50<sup>a</sup>±1.00 | 30.25<sup>b</sup>±0.59 |
| 6. | Eosonophil% | 16. 62<sup>a</sup>±0.49 | 2.12<sup>b</sup> ±0.44 |
| 7. | Basophil% | 0.00 | 0.00 |
| 8. | Lymphocyte% | 29.12<sup>b</sup>±0.29 | 60.87<sup>a</sup>±0.29 |
| 9. | Monocyte% | 1.37±0.18 | 1.87±0.29 |

*The values that have superscripts was showing significant differences between groups. The superscript 'a' was showing high mean value than 'b'.Significant at (p ≤0.05).*

In healthy and diseased sheep, respectively, the mean hemoglobin values were determined to be 10.37^a^±0.18344 and 7.96^b^±0.19329gm/dl, as indicated in **Table 1**. In the current investigation, a diseased sheep’s hemoglobin level was significantly lower than that of a healthy sheep (p ≤0.05). The current finding was consistent with previously reported results **(Ved Prakash et al., 2010; Shekh I. J et al., 2017; Tiwari A.N. et al., 2017)**. The L4 larvae and adult worms drain blood from the animal’s abomasum, causing anemia and edoema, which may be the cause of the value decline. According to reports, H. contortus infection causes an average blood loss of 0.05 ml per day per worm **(Urquhart et al., 2000).**In healthy and diseased sheep, respectively, the mean ESR values were 0.9063±0.04575 and 1.0000 ±0.00 mm**(Table1)**. Non-significant differences between diseased and healthy sheep are revealed by the analysis of variance for ESR (p ≤0.05).ESR did not significantly alter in the current research **(Tiwari et al.,2017).**While mean PCV values of 36.2550^a^±0.6422 and 27.8425^b^±0.67929 % were observed in healthy and diseased sheeprespectively**(Table 1).** In the current investigation, diseased sheep had significantly lower PCV levels than healthy sheep. The current findings were consistent with the previous findings**(Ved Prakash et al., 2010; Shekh I. J et al., 2018; Tiwari A.N. et al., 2017).** The drop in PCV levels in infected sheep may be caused by bleeding from the abomasa as a result of parasite-induced wounds and by a delay between blood loss and the activation of the erythropoietic system to make up for blood loss **(Dargie and Allonby 1975).**In the current study, a drop in TEC values (p ≤0.05)in infected sheep may be caused by bleeding from the abdomen due to wounds created by the H. contortus, as well as by a delay between blood loss and the activation of the erythropoietic system to make up for blood loss **(Dargie and Allonby 1975).**Neutrophilia and eosinophilia may be responsible for the diseased sheep’s higher total leukocyte count than the healthy ones **(Sheikh et al.,2018).** In comparison to healthy sheep, a significant rise in neutrophil counts was seen in diseased sheep. As worm burden grows, neutrophil levels rise. It is recognized as a significant chemotaxis parasiticide effector and as a marker of efficient host defense against helminthiasis **(Okoye et al., 2013).**When compared to healthy sheep, a substantial rise in eosinophil count was found. An essential sign of helminth infection is an increase in blood eosinophils **(Soulsby, 1982).** Eosinophils and antibodies work together to combat parasites and assist the host get rid of them **(Parsani et al., 2011).**Significantly fewer lymphocytes were seen in the diseased sheep in the current investigation. The reduction in lymphocytes may be brought on by the infiltration of these cells into other organs and the killing of lymphocytes in lymphoid organs **(Sandhu et al., 1998).** Similar results were observed in a study of sheep helminth infection **(Sheikh et al.,2018; Okoye et al.,2013).**Between healthy and diseased sheep, there were no discernible differences in basophil values, however, there were non-significant differences in monocyte values.Overall, significant differences were observed in Hb gm/dl, PCV%, Neutrophil %, Eosinophil %, TEC /cumm, TLC/cumm, and Lymphocyte %. Better hemoglobin, PCV, and TEC was observed in healthy sheep in comparison to that diseased. However, total leucocyte count was pronounced in diseased (infected) sheep. Eosinophil count was found to have significantly increased in infected sheep. Both neutrophils and lymphocytes showed a similarly substantial rise, indicating immunological response was involved.

The comparative biochemical parameters are illustrated in **Table 2**.

**Table 2:** biochemical parameters (liver function test)in diseased and healthy sheep.

| Sr. no | Parameters | Diseased | Healthy |
| --- | --- | --- | --- |
| 1 | Total protein (g/dl) | 6.11<sup>b</sup>±0.02 | 7.77<sup>a</sup>±0.03 |
| 2 | Albumin (g/dl) | 1.92<sup>b</sup>±0.04 | 2.80<sup>a</sup>±0.03 |
| 3 | Globulin(g/dl) | 4.18<sup>a</sup>±0.06 | 4.97<sup>b</sup>±0.06 |
| 4 | Albumin:Globulin | 0.46<sup>b</sup>±0.01 | 0.58<sup>a</sup>±0.01 |
| 5. | S.G.P.T IU/L | 39.75<sup>a</sup>±1.4 | 32.12<sup>b</sup>±1.0 |
| 6 | S.G.O.T IU/L | 86.87<sup>a</sup>±2.6 | 68.87<sup>b</sup>±2.7 |
| 7 | Alkaline Phosphatase IU/L | 85.25<sup>a</sup>±3.89 | 64.87<sup>b</sup>±2.09 |
| 8 | Total Bilirubin mg/dl | 0.67<sup>a</sup>±0.07 | 0.37<sup>b</sup>±0.03 |
| 9 | Direct Bilirubin mg/dl | 0.30<sup>a</sup>±0.02 | 0.16<sup>b</sup>±0.01 |
| 10 | Indirect Bilirubin mg/dl | 0.37<sup>a</sup>±0.09014 | 0.21<sup>b</sup>±0.05 |

*The Mean values that have a superscript, was showing significant differences between groups. The superscript 'a' was showing high mean value than 'b'.Significant at (p ≤0.05).*

When compared to healthy sheep, there was a significant decrease in total protein levels. The current findings are consistent with those obtained in goats by **Purohit et al. (2003), Ashok Arora et al. (2003).** Since HaemonchusControtus is a blood-sucking parasite and damages intestinal mucosa, which results in inappropriate digestion and poor protein absorption, the lower amount of total protein in the infected animal may be caused by increased serum leakage **(Radostits et al., 1994).** Additionally, it was observed that an animal infected with Haemonchus had a mean daily feces clearance of 210–340 ml/day **(Dargie and Allonby, 1975).**Between healthy and diseased sheep in the current investigation, albumin levels were significantly reduced, as shown in (Table 2). A high value was observed in healthy sheep than in disease sheep. The current findings were consistent with those**(Ashok Kumar et al., 2005; Arora et al. 2003; Nicholson et al., 2000).** According to **Tanwar and Mishra (2001),** hypoalbuminemia may result from the selective loss of albumin that is smaller in size and more osmotically sensitive to fluid flow. It may also result from enhanced albumin catabolism and protein malabsorption owing to the damaged intestinal mucosa **(Catchpole and Gregory, 1985).**

In the current investigation, it was shown that diseased sheep had much lower levels of globulin than healthy sheep did. According to **(Dhanlakshmi et al., 2002),** the major cause of hypoproteinemia may be inappetence, which leads to a decrease in dietary protein intake and plasma losses from damaged intestinal mucosa **(Purohit et al., 2003).** In healthy and diseased sheep, there are appreciable variations in the A:G ratio (p≤ 0.05). In the current investigation, diseased sheep had significantly lower levels of the A/G ratio than healthy sheep. The current findings were consistent with those**(Maiti et al.1999; Pandit et al.2009).** According to research, changes in albumin and globulin levels led to a decrease in the A/G ratio **(Hosseini et al., 2012).**In the current investigation, diseased sheep had significantly higher levels of total bilirubin than healthy sheep. The current findings were consistent with those**(Zaki et al. 2003; Kumar et al. 2015).** Infected animals had much higher levels of direct bilirubin than healthy sheep. As demonstrated, there is a non-significant difference in indirect bilirubin levels between healthy and ill sheep **(Table 2).** The current findings are consistent with **(Zaki et al. 2003).** Hepatocyte injury and intestinal pathology might both contribute to an increase in bilirubin **(Radostits et al.,1994).**Infected animals had significantly higher levels of SGPT and SGOT than healthy sheep. The current finding was consistent with **(Prasanthi et al.1999; Zaki et al., 2003).** According to **Sharma and Joshi (2001),** in sheep infected with *H. contortus*, the large elevations in SGPT and SGOT levels may be caused by increased membrane permeability and abnormalities in hepatic function.When compared to healthy sheep, diseased animals had a much higher level of alkaline phosphatase. An increase in the alkaline phosphatase value may be caused by damaged intestinal mucosal cells as a result of parasitic pathogenesis in GIT disorders because alkaline phosphatase is widely distributed throughout the body and is concentrated in bone, intestinal mucosa, renal tubules cells, liver, and placenta **(Tiwari et al., 2017).**Overall, the Total protein, globulin, albumin, and albumin: globulin ratio was observed to be better in healthy sheep. While other reports indicate more in infected sheep (S.G.P.T. S.G.O.T, Total Bilirubin, Alkaline Phosphatase, Direct Bilirubin, Indirect Bilirubin).

Comparative biochemical parameters are presented in **Table 3**.

**Table 3.** Biochemical parameters (kidney function test) in diseased and healthy sheep.

| S.no | Parameter | Diseases | Healthy |
| --- | --- | --- | --- |
| 1 | Glucose mg/dl | 55.50<sup>b</sup>±1.21 | 68.62<sup>a</sup>±2.33 |
| 2 | Creatinine mg/dl | 1.71<sup>a</sup>±0.05 | 1.20<sup>b</sup>±0.02 |
| 3 | Uric acid mg/dl | 0.89<sup>a</sup>±0.01 | 0.76<sup>b</sup>±0.04311 |
| 4 | Urea mg/dl | 48.37<sup>a</sup>±1.2 | 36.12<sup>b</sup>±1.56 |
| 5 | BUN mg/dl | 23.62<sup>a</sup>±0.73 | 16.87<sup>b</sup>±0.95 |

## Discussion

The current study aims to explore the host immunity against dreadly gastrointestinal parasite Haemonchus contortus in sheep model. Although this is a less explored field, we have earlier documented RIGI (well known antiviral molecule) and CD14 (well known antibacterial molecule) to have a potent role in providing host immunity against *Haemonchus contortus* (Banerjee et al., 2021, Rawat et al., 2021). In this study, we had explored IL6 and IL10 as potent immune response genes for host immunity against Haemonchus contortus. We had characterized IL6 and IL10 genes for Garole sheep for the first time. Certain sites for post translational modification have been detected in both IL6 and IL10 of Garole sheep. These are sites for N-linked glycosylation, aids in solubilization in water. Others are site for di sulphide linkage, that are helpful for providing appropriate 3D structure to the molecule. We had also depicted the 3D structure of these molecules. Molecular docking analysis have revealed that IL10 act as receptor for Haemochus contortus. It binds with the surface membrane for H. contortus and the binding sites are being predicted. Although scanty reports are available for the role of these immune response molecules against parasitic infestation, such as chagus disease, trypanosomiasis, toxoplasmosis and documented through KEGG analysis. However, this is the first report for role of these molecules in host resistance against H. contortus. One interesting observation is that host resistance against pathogen is governed by a cascade of immunological processes mediated through a number of molecules revealed through STRING analysis. Hence resistance trait is polygenic in inheritance.

We have sequenced both IL10 and IL6 molecule and characterized them through a series of in silico approaches, including molecular docking. In the next step we have validated them through differential mRNA expression profiling, haematological and biochemical estimation, followed by confirmation through immunohistochemistry.

Earlier report for characterization of these genes in swamp buffalo indicate similar result. Swamp buffalo was found to have a similar amino acid sequence **(Mingala et al., 2007).** From 1 to 25 aa, a signal peptide was anticipated. 27 sites were identified for O-linked glycosylation sites. At positions 72–78 and 101–111, there are 4 and 2 disulphide bonds between cysteines, respectively. Similar to this, one N-linked glycosylation site was discovered in the IL-6 gene of the swamp buffalo, and four cysteine residues are crucial in defining the tertiary structure and functional integrity of the protein**(Mingala et al., 2007).** acetylation site was detected (MNSLFT). The stability, location, synthesis, and interactions of proteins are all influenced by the N-terminal acetylation **(Hollebeke et al., 2012)**.The C-mannosylation site was reported to start at amino acid position 181. In supporting cell-to-cell adhesion and communication, it plays a critical biological role **(Loke et al., 2016).** A leucine-rich nuclear export signal was not found at any location. 20 protein phosphorylation sites were found at the following amino acid positions: 3, 6, 7, 10, 15, 28, 41, 43, 48, 49, 65, 81, 84, 116, 139, 145, 140, 149, 193, 201. In signalling pathways and metabolism, phosphorylation plays a significant and well-established function**(Cie**ś**la et al., 2011).**Two domains (1-81, 82-208) were predicted by the protein sequence analysis. The protein domain performs a lot of functions, such as binding a specific molecule or catalyzing a specific process, which together contributes to the protein’s overall activity **(Wetlaufer, 1973).** Protein sequences in the regions 17-53, 25-42, 41-52, and 159-167 included the domain linker. Domain linkers are sequences that connect two structural domains and stop folding domains from interfering unfavorably**(Tanaka et al., 2003).**At 18 and 165, transmembrane helices are located. Transmembrane proteins are essential membrane components that span the whole-cell membrane to which they are permanently connected. Numerous transmembrane proteins serve as passages that permit the movement of certain molecules through the membrane. To transport a drug through the membrane, they usually go through major conformational changes. Alpha-helical transmembrane proteins, such as potassium ions, are voltage-gated ion channels **(Purves et al., 2001).**At the c terminal’s sequence position 177, a GPI modification site was identified. It performs a variety of biological processes, including cell-cell adhesion and interaction **(Low,1999).** The GPI anchor may also serve as a bridge connecting a cell’s outside and its interior signaling molecules (Robinson, 1997). **Figure 2A** illustrates the results of secondary structure and disordered protein prediction (**Figure 2B)**, which showed that 70% of the protein was an alpha helix, 0% was a beta-strand, and 44% was a disordered protein. **Figure 3** illustrates the helical shape of the IL-6 gene, which is represented as the structure of the IL-6 gene using Pymol.

Molecular docking analysis revealed that IL10 act as a receptor molecule for H. contortus. Earlier studies have also reported that IL10 too act as receptor. In our earlier studies while working with RIGI and CD14 as receptor and prediction of binding site, we have worked with molecular docking (Banerjee etal., 2021, Rawat et al.,2021).

IL-6 is a pleiotropic cytokine that acts in both a pro- and anti-inflammatory manner to affect a number of functions, including immunology, tissue repair, and metabolism**(Scheller et al.,2011).**Similar to other pro-inflammatory cytokines like TNF-alpha and IL-6, IL-1β, is an endogenous pyrogen that causes fever and the generation of acute-phase proteins from the liver.Additionally, it promotes B cell maturation into plasma cells andstimulates cytotoxic T cells**(Barkhausen et al., 2011).**In the current study, the IL-6 gene’s mRNA expression was lower in diseased sheep than in healthy ones. Studies that detailed immune regulation mechanisms against helminths demonstrate, in contrast to our findings, that in a tropical environment, differences in the expression of the IL-5 and IL-6 genes in Pelibuey sheep can affect the phenotypic traits that confer resistance to and susceptibility to H. contortus **(Estrada-Reyes et al.,2015).**The abomasal tissue of resistant Angus heifers infected with the GINs Cooperiaoncophora and Ostertagiaostertagi produced higher IL-6 and other proinflammatory cytokines, according to studies of a similar kind**(Lie et al. 2007)**. According to the current research, diseased sheep have higher expression of the IL-10 gene and lower expression of the IL-6 gene. Given that IL-10 is an immunosuppressive and anti-inflammatory cytokine that controls inflammation as well as T cell, NK cell, and macrophage activity, decreased expression of the IL-6 gene may be caused by increased expression of the IL-10 gene **(Kamnaka et al., 2006)**. By preventing CD28 phosphorylation, IL-10 specifically prevents co-stimulation of T cells, which prevents the production of Th1 cytokines like IL-6 **(Moore et al.,2001; Tizard, 2004)**.

In the current investigation, a diseased sheep’s hemoglobin level was significantly lower than that of a healthy sheep (p ≤0.05). The current finding was consistent with previously reported results **(Ved Prakash et al., 2010; Shekh I. J et al., 2017; Tiwari A.N. et al., 2017)**. The L4 larvae and adult worms drain blood from the animal’s abomasum, causing anemia and oedema, which may be the cause of the value decline. According to reports, H. contortus infection causes an average blood loss of 0.05 ml per day per worm **(Urquhart et al., 2000).** In healthy and diseased sheep, respectively, the mean ESR values were 0.9063±0.04575 and 1.0000 ±0.00 mm

## Discussion

The current study aims to explore the host immunity against dreadly gastrointestinal parasite Haemonchus contortus in sheep model. Although this is a less explored field, we have earlier documented RIGI (well known antiviral molecule) and CD14 (well known antibacterial molecule) to have a potent role in providing host immunity against Haemonchus contortus (Banerjee *et al*, 2021, Rawat *et al*., 2021). In this study, we had explored IL6 and IL10 as potent immune response genes for host immunity against Haemonchus contortus. We had characterized IL6 and IL10 genes for Garole sheep for the first time. Certain sites for post translational modification have been detected in both IL6 and IL10 of Garole sheep. These are sites for N-linked glycosylation, aids in solubilization in water. Others are site for di sulphide linkage, that are helpful for providing appropriate 3D structure to the molecule. We had also depicted the 3D structure of these molecules. Molecular docking analysis have revealed that IL10 act as receptor for Haemochus contortus. It binds with the surface membrane for H. contortus and the binding sites are being predicted. Although scanty reports are available for the role of these immune response molecules against parasitic infestation, such as chagus disease, trypanosomiasis, toxoplasmosis and documented through KEGG analysis . However, this is the first report for role of these molecules in host resistance against H. contortus. One interesting observation is that host resistance against pathogen is governed by a cascade of immunological processes mediated through a number of molecules revealed through STRING analysis. Hence resistance trait is polygenic in inheritance.

We have sequenced both IL10 and IL6 molecule and characterized them through a series of in silico approaches, including molecular docking. In the next step we have validated them through differential mRNA expression profiling, haematological and biochemical estimation, followed by confirmation through immunohistochemistry.

Earlier report for characterization of these genes in swamp buffalo indicate similar result. Swamp buffalo was found to have a similar amino acid sequence **(Mingala et al., 2007).**From 1 to 25 aa, a signal peptide was anticipated. 27 sites were identified for O-linked glycosylation sites. At positions 72–78 and 101–111, there are 4 and 2 disulphide bonds between cysteines, respectively. Similar to this, one N-linked glycosylation site was discovered in the IL-6 gene of the swamp buffalo, and four cysteine residues are crucial in defining the tertiary structure and functional integrity of the protein**(Mingala et al., 2007).** acetylation site was detected (MNSLFT). The stability, location, synthesis, and interactions of proteins are all influenced by the N-terminal acetylation **(Hollebeke et al., 2012)**.The C-mannosylation site was reported to start at amino acid position 181. In supporting cell-to-cell adhesion and communication, it plays a critical biological role**(Loke et al., 2016).** A leucine-rich nuclear export signal was not found at any location. 20 protein phosphorylation sites were found at the following amino acid positions: 3, 6, 7, 10, 15, 28, 41, 43, 48, 49, 65, 81, 84, 116, 139, 145, 140, 149, 193, 201. In signalling pathways and metabolism, phosphorylation plays a significant and well-established function**(Cie**ś**la et al., 2011).**2 domains (1-81, 82-208) were predicted by the protein sequence analysis. The protein domain performs a lot of functions, such as binding a specific molecule or catalyzing a specific process, which together contributes to the protein’s overall activity **(Wetlaufer, 1973).** Protein sequences in the regions 17-53, 25-42, 41-52, and 159-167 included the domain linker. Domain linkers are sequences that connect two structural domains and stop folding domains from interfering unfavorably**(Tanaka et al., 2003).**At 18 and 165, transmembrane helices are located. Transmembrane proteins are essential membrane components that span the whole-cell membrane to which they are permanently connected. Numerous transmembrane proteins serve as passages that permit the movement of certain molecules through the membrane. To transport a drug through the membrane, they usually go through major conformational changes. Alpha-helical transmembrane proteins, such as potassium ions, are voltage-gated ion channels **(Purves et al., 2001).**At the c terminal’s sequence position 177, a GPI modification site was identified. It performs a variety of biological processes, including cell-cell adhesion and interaction **(Low,1999).** The GPI anchor may also serve as a bridge connecting a cell’s outside and its interior signaling molecules (Robinson, 1997). **Figure 2A** illustrates the results of secondary structure and disordered protein prediction (**Figure 2B)**, which showed that 70% of the protein was an alpha helix, 0% was a beta-strand, and 44% was a disordered protein. **Figure 3** illustrates the helical shape of the IL-6 gene, which is represented as the structure of the IL-6 gene using Pymol.

Molecular docking analysis revealed that IL10 act as a receptor molecule for H. contortus. Earlier studies have also reported that IL10 too act as receptor. In our earlier studies while working with RIGI and CD14 as receptor and prediction of binding site, we have worked with molecular docking (Banerjee et al., 2021, Rawat et al., 2021).

IL-6 is a pleiotropic cytokine that acts in both a pro- and anti-inflammatory manner to affect a number of functions, including immunology, tissue repair, and metabolism**(Scheller et al.,2011).**Similar to other pro-inflammatory cytokines like TNF-alpha and IL-6, IL-1β, is an endogenous pyrogen that causes fever and the generation of acute-phase proteins from the liver.Additionally, it promotes B cell maturation into plasma cells andstimulates cytotoxic T cells**(Barkhausen et al., 2011).**In the current study, the IL-6 gene’s mRNA expression was lower in diseased sheep than in healthy ones. Studies that detailed immune regulation mechanisms against helminths demonstrate, in contrast to our findings, that in a tropical environment, differences in the expression of the IL-5 and IL-6 genes in Pelibuey sheep can affect the phenotypic traits that confer resistance to and susceptibility to H. contortus**(Estrada-Reyes et al.,2015).**The abomasal tissue of resistant Angus heifers infected with the GINs Cooperiaoncophora and Ostertagiaostertagi produced higher IL-6 and other proinflammatory cytokines, according to studies of a similar kind**(Lie et al. 2007)**. According to the current research, diseased sheep have higher expression of the IL-10 gene and lower expression of the IL-6 gene. Given that IL-10 is an immunosuppressive and anti-inflammatory cytokine that controls inflammation as well as T cell, NK cell, and macrophage activity, decreased expression of the IL-6 gene may be caused by increased expression of the IL-10 gene **(Kamnaka et al., 2006)**. By preventing CD28 phosphorylation, IL-10 specifically prevents co-stimulation of T cells, which prevents the production of Th1 cytokines like IL-6 **(Moore et al.,2001; Tizard, 2004)**.

In the current investigation, a diseased sheep’s hemoglobin level was significantly lower than that of a healthy sheep (p ≤0.05). The current finding was consistent with previously reported results **(Ved Prakash et al., 2010; Shekh I. J et al., 2017; Tiwari A.N. et al., 2017)**. The L4 larvae and adult worms drain blood from the animal’s abomasum, causing anemia and edoema, which may be the cause of the value decline. According to reports, H. contortus infection causes an average blood loss of 0.05 ml per day per worm **(Urquhart et al., 2000).**In healthy and diseased sheep, respectively, the mean ESR values were 0.9063±0.04575 and 1.0000 ±0.00 mm**(Table1)**. Non-significant differences between diseased and healthy sheep are revealed by the analysis of variance for ESR (p ≤0.05).ESR did not significantly alter in the current research **(Tiwari et al.,2017).**While mean PCV values of 36.2550^a^±0.6422 and 27.8425^b^±0.67929 % were observed in healthy and diseased sheeprespectively**(Table 1).** In the current investigation, diseased sheep had significantly lower PCV levels than healthy sheep. The current findings were consistent with the previous findings**(Ved Prakash et al., 2010; Shekh I. J et al., 2018; Tiwari A.N. et al., 2017).** The drop in PCV levels in infected sheep may be caused by bleeding from the abomasa as a result of parasite-induced wounds and by a delay between blood loss and the activation of the erythropoietic system to make up for blood loss **(Dargie and Allonby 1975).**In the current study, a drop in TEC values (p ≤0.05)in infected sheep may be caused by bleeding from the abdomen due to wounds created by the H. contortus, as well as by a delay between blood loss and the activation of the erythropoietic system to make up for blood loss **(Dargie and Allonby 1975).**Neutrophilia and eosinophilia may be responsible for the diseased sheep’s higher total leukocyte count than the healthy ones **(Sheikh et al.,2018).** In comparison to healthy sheep, a significant rise in neutrophil counts was seen in diseased sheep. As worm burden grows, neutrophil levels rise. It is recognized as a significant chemotaxis parasiticide effector and as a marker of efficient host defense against helminthiasis **(Okoye et al., 2013).**When compared to healthy sheep, a substantial rise in eosinophil count was found. An essential sign of helminth infection is an increase in blood eosinophils **(Soulsby, 1982).** Eosinophils and antibodies work together to combat parasites and assist the host get rid of them **(Parsani et al., 2011).**Significantly fewer lymphocytes were seen in the diseased sheep in the current investigation. The reduction in lymphocytes may be brought on by the infiltration of these cells into other organs and the killing of lymphocytes in lymphoid organs **(Sandhu et al., 1998).** Similar results were observed in a study of sheep helminth infection **(Sheikh et al.,2018;Okoye et al.,2013).**Between healthy and diseased sheep, there were no discernible differences in basophil values, however, there were non-significant differences in monocyte values.Overall, significant differences were observed in Hb gm/dl, PCV%, Neutrophil %, Eosinophil %, TEC /cumm, TLC/cumm, and Lymphocyte %. Better hemoglobin, PCV, and TEC was observed in healthy sheep in comparison to that diseased. However, total leucocyte count was pronounced in diseased (infected) sheep. Eosinophil count was found to have significantly increased in infected sheep. Both neutrophils and lymphocytes showed a similarly substantial rise, indicating immunological response was involved.

When compared to healthy sheep, there was a significant decrease in total protein levels. The current findings are consistent with those obtained in goats by **Purohit et al. (2003), Ashok Arora et al. (2003).** Since HaemonchusControtus is a blood-sucking parasite and damages intestinal mucosa, which results in inappropriate digestion and poor protein absorption, the lower amount of total protein in the infected animal may be caused by increased serum leakage **(Radostits et al., 1994).** Additionally, it was observed that an animal infected with Haemonchus had a mean daily feces clearance of 210–340 ml/day **(Dargie and Allonby, 1975).**Between healthy and diseased sheep in the current investigation, albumin levels were significantly reduced, as shown in (Table 2). A high value was observed in healthy sheep than in disease sheep. The current findings were consistent with those**(Ashok Kumar et al., 2005; Arora et al. 2003; Nicholson et al., 2000).** According to **Tanwar and Mishra (2001),** hypoalbuminemia may result from the selective loss of albumin that is smaller in size and more osmotically sensitive to fluid flow. It may also result from enhanced albumin catabolism and protein malabsorption owing to the damaged intestinal mucosa **(Catchpole and Gregory, 1985).**

In the current investigation, it was shown that diseased sheep had much lower levels of globulin than healthy sheep did. According to **(Dhanlakshmi et al., 2002),** the major cause of hypoproteinemia may be inappetence, which leads to a decrease in dietary protein intake and plasma losses from damaged intestinal mucosa **(Purohit et al., 2003).** In healthy and diseased sheep, there are appreciable variations in the A:G ratio (p≤ 0.05). In the current investigation, diseased sheep had significantly lower levels of the A/G ratio than healthy sheep. The current findings were consistent with those**(Maiti et al.1999; Pandit et al.2009).** According to research, changes in albumin and globulin levels led to a decrease in the A/G ratio **(Hosseini et al., 2012).**In the current investigation, diseased sheep had significantly higher levels of total bilirubin than healthy sheep. The current findings were consistent with those**(Zaki et al. 2003; Kumar et al. 2015).** Infected animals had much higher levels of direct bilirubin than healthy sheep. As demonstrated, there is a non-significant difference in indirect bilirubin levels between healthy and ill sheep **(Table 2).** The current findings are consistent with **(Zaki et al. 2003).** Hepatocyte injury and intestinal pathology might both contribute to an increase in bilirubin **(Radostits et al.,1994).**Infected animals had significantly higher levels of SGPT and SGOT than healthy sheep. The current finding was consistent with **(Prasanthi et al.1999; Zaki et al., 2003).** According to **Sharma and Joshi (2001),** in sheep infected with *H. contortus*, the large elevations in SGPT and SGOT levels may be caused by increased membrane permeability and abnormalities in hepatic function.When compared to healthy sheep, diseased animals had a much higher level of alkaline phosphatase. An increase in the alkaline phosphatase value may be caused by damaged intestinal mucosal cells as a result of parasitic pathogenesis in GIT disorders because alkaline phosphatase is widely distributed throughout the body and is concentrated in bone, intestinal mucosa, renal tubules cells, liver, and placenta **(Tiwari et al., 2017).**Overall, the Total protein, globulin, albumin, and albumin: globulin ratio was observed to be better in healthy sheep. While other reports indicate more in infected sheep (S.G.P.T. S.G.O.T, Total Bilirubin, Alkaline Phosphatase, Direct Bilirubin, Indirect Bilirubin).

The glucose level in diseased sheep was much lower in our study than in healthy sheep, which was consistent with **(Ashok Kumar et al., 2005)** who found low glucose levels in goats infected with Haemonchus species. Significantly lower food intake and poor nutrient digestion brought on by parasite illnesses’ gastrointestinal problems may also contribute to significant reductions in glucose levels **(Hayat et al., 1996).** Reduced bloodstream absorption and quick digestion and uptake of soluble fats and carbohydrates from the stomach by parasites **(Pathak and Tiwari 2012).** Diseased sheep show a significant increase in serum urea and uric acid levels. Impaired regulation of renal tubular transport may be the cause of the increased value of these two measures in sick sheep compared to healthy sheep **(Hiranyachattada et al., 2000).** Significant increases in serum BUN and urea levels in sick sheep compared to healthy sheep may be caused by liver damage from parasite infection and decreased control of renal tubular transport **(Hiranyachattada et al., 2000; Yousif et al., 1990).**

. Non-significant differences between diseased and healthy sheep are revealed by the analysis of variance for ESR (p ≤0.05).ESR did not significantly alter in the current research **(Tiwari et al.,2017).**While mean PCV values of 36.2550^a^±0.6422 and 27.8425^b^±0.67929 % were observed in healthy and diseased sheeprespectively**(Table 1).** In the current investigation, diseased sheep had significantly lower PCV levels than healthy sheep. The current findings were consistent with the previous findings**(Ved Prakash et al., 2010; Shekh I. J et al., 2018; Tiwari A.N. et al., 2017).** The drop in PCV levels in infected sheep may be caused by bleeding from the abomasa as a result of parasite-induced wounds and by a delay between blood loss and the activation of the erythropoietic system to make up for blood loss **(Dargie and Allonby 1975).**In the current study, a drop in TEC values (p ≤0.05)in infected sheep may be caused by bleeding from the abdomen due to wounds created by the H. contortus, as well as by a delay between blood loss and the activation of the erythropoietic system to make up for blood loss **(Dargie and Allonby 1975).**Neutrophilia and eosinophilia may be responsible for the diseased sheep’s higher total leukocyte count than the healthy ones **(Sheikh et al.,2018).** In comparison to healthy sheep, a significant rise in neutrophil counts was seen in diseased sheep. As worm burden grows, neutrophil levels rise. It is recognized as a significant chemotaxis parasiticide effector and as a marker of efficient host defense against helminthiasis **(Okoye et al., 2013).**When compared to healthy sheep, a substantial rise in eosinophil count was found. An essential sign of helminth infection is an increase in blood eosinophils **(Soulsby, 1982).** Eosinophils and antibodies work together to combat parasites and assist the host get rid of them **(Parsani et al., 2011).**Significantly fewer lymphocytes were seen in the diseased sheep in the current investigation. The reduction in lymphocytes may be brought on by the infiltration of these cells into other organs and the killing of lymphocytes in lymphoid organs **(Sandhu et al., 1998).** Similar results were observed in a study of sheep helminth infection **(Sheikh et al.,2018;Okoye et al.,2013).**Between healthy and diseased sheep, there were no discernible differences in basophil values, however, there were non-significant differences in monocyte values.Overall, significant differences were observed in Hb gm/dl, PCV%, Neutrophil %, Eosinophil %, TEC /cumm, TLC/cumm, and Lymphocyte %. Better hemoglobin, PCV, and TEC was observed in healthy sheep in comparison to that diseased. However, total leucocyte count was pronounced in diseased (infected) sheep. Eosinophil count was found to have significantly increased in infected sheep. Both neutrophils and lymphocytes showed a similarly substantial rise, indicating immunological response was involved.

When compared to healthy sheep, there was a significant decrease in total protein levels. The current findings are consistent with those obtained in goats by **Purohit et al. (2003), Ashok Arora et al. (2003).** Since HaemonchusControtus is a blood-sucking parasite and damages intestinal mucosa, which results in inappropriate digestion and poor protein absorption, the lower amount of total protein in the infected animal may be caused by increased serum leakage **(Radostits et al., 1994).** Additionally, it was observed that an animal infected with Haemonchus had a mean daily feces clearance of 210–340 ml/day **(Dargie and Allonby, 1975).**Between healthy and diseased sheep in the current investigation, albumin levels were significantly reduced, as shown in (Table 2). A high value was observed in healthy sheep than in disease sheep. The current findings were consistent with those **(Ashok Kumar et al., 2005; Arora et al. 2003; Nicholson et al., 2000).** According to **Tanwar and Mishra (2001),** hypoalbuminemia may result from the selective loss of albumin that is smaller in size and more osmotically sensitive to fluid flow. It may also result from enhanced albumin catabolism and protein malabsorption owing to the damaged intestinal mucosa **(Catchpole and Gregory, 1985).**

In the current investigation, it was shown that diseased sheep had much lower levels of globulin than healthy sheep did. According to **(Dhanlakshmi et al., 2002),** the major cause of hypoproteinemia may be inappetence, which leads to a decrease in dietary protein intake and plasma losses from damaged intestinal mucosa **(Purohit et al., 2003).** In healthy and diseased sheep, there are appreciable variations in the A:G ratio (p≤ 0.05). In the current investigation, diseased sheep had significantly lower levels of the A/G ratio than healthy sheep. The current findings were consistent with those**(Maiti et al.1999; Pandit et al.2009).** According to research, changes in albumin and globulin levels led to a decrease in the A/G ratio **(Hosseini et al., 2012).**In the current investigation, diseased sheep had significantly higher levels of total bilirubin than healthy sheep. The current findings were consistent with those**(Zaki et al. 2003; Kumar et al. 2015).** Infected animals had much higher levels of direct bilirubin than healthy sheep. As demonstrated, there is a non-significant difference in indirect bilirubin levels between healthy and ill sheep **(Table 2).** The current findings are consistent with **(Zaki et al. 2003).** Hepatocyte injury and intestinal pathology might both contribute to an increase in bilirubin **(Radostits et al.,1994).**Infected animals had significantly higher levels of SGPT and SGOT than healthy sheep. The current finding was consistent with **(Prasanthi et al.1999; Zaki et al., 2003).** According to **Sharma and Joshi (2001),** in sheep infected with *H. contortus*, the large elevations in SGPT and SGOT levels may be caused by increased membrane permeability and abnormalities in hepatic function.When compared to healthy sheep, diseased animals had a much higher level of alkaline phosphatase. An increase in the alkaline phosphatase value may be caused by damaged intestinal mucosal cells as a result of parasitic pathogenesis in GIT disorders because alkaline phosphatase is widely distributed throughout the body and is concentrated in bone, intestinal mucosa, renal tubules cells, liver, and placenta **(Tiwari et al., 2017).**Overall, the Total protein, globulin, albumin, and albumin: globulin ratio was observed to be better in healthy sheep. While other reports indicate more in infected sheep (S.G.P.T. S.G.O.T, Total Bilirubin, Alkaline Phosphatase, Direct Bilirubin, Indirect Bilirubin).

The glucose level in diseased sheep was much lower in our study than in healthy sheep, which was consistent with **(Ashok Kumar et al., 2005)** who found low glucose levels in goats infected with Haemonchus species. Significantly lower food intake and poor nutrient digestion brought on by parasite illnesses’ gastrointestinal problems may also contribute to significant reductions in glucose levels **(Hayat et al., 1996).** Reduced bloodstream absorption and quick digestion and uptake of soluble fats and carbohydrates from the stomach by parasites **(Pathak and Tiwari 2012).** Diseased sheep show a significant increase in serum urea and uric acid levels. Impaired regulation of renal tubular transport may be the cause of the increased value of these two measures in sick sheep compared to healthy sheep **(Hiranyachattada et al., 2000).** Significant increases in serum BUN and urea levels in sick sheep compared to healthy sheep may be caused by liver damage from parasite infection and decreased control of renal tubular transport **(Hiranyachattada et al., 2000; Yousif et al., 1990).**

## Conclusion

In conclusion, this study describes the molecular characterization of IL-6 in sheep, which offers fundamental information required to advance the study of functional immune responses in this animal, may help understand the antibacterial and antiparasitic immunity in sheep, enables a more in-depth examination of these sheep’s immune responses, and presents the possibility of using them as recombinant proteins to manipulate the immune response. Now that IL-6 and IL10 is being produced, it may be possible to evaluate its value in vaccine research and to take a broader view of its function in the immune response.The differential mRNA expression level of the immune response gene in healthy and disease sheep described that the differentially expressed gene IL-6 has a major role in homeostasis maintenance and immune responses.The immune-responsive gene discovered here offers fresh research avenues for the investigation of sheep long-term resistance to H. contortus infection.

